# Long-term stress shapes dynamic reconfiguration of functional brain networks across multi-task demands

**DOI:** 10.1101/2023.03.28.534193

**Authors:** Hongyao Gao, Yimeng Zeng, Ting Tian, Chao Liu, Jianhui Wu, Haitao Wu, Shaozheng Qin

## Abstract

Exposure to sustained stress can have a profound impact on the brain, emotion and cognition, with either adaptive or maladaptive effects. Human functional brain networks are dynamically organized to enable rapid and flexible adaptation to meet ever-changing task demands. Yet, little is known about how long-term stress alters the dynamic reconfiguration of functional brain networks across multi-task demands. Here we show prominent changes in the dynamic reconfiguration of large-scale brain networks during resting-state, emotional and working-memory processing under long-term stress. Hidden Markov Model analysis detected several latent brain states and switching processes involving the default mode, emotional salience and executive-control networks that are dominant to rest, emotion and working memory, respectively. Critically, long-term stress increased persistent time on brain states relevant to goal-directed demands and cognitive control, with more frequent transitions to these brain states when compared to controls. Furthermore, long-term stress led to higher correlations of the occupancy and persistency of brain states linked to psychological distress and behavioral performance. Our findings provide a neurocognitive framework whereby long-term stress shapes the way the brain adapts to varying task demands and increases the sensitivity of functional brain networks to psychological and behavioral responses. These changes can be both adaptive and maladaptive, reflecting the complex effects of long-term stress on brain function.

## Introduction

The human brain’s ability to rapidly and flexibly adapt to ever-changing environmental and task demands is a hallmark of intelligence. Neuroimaging research has demonstrated that functional brain networks are dynamically organized into certain manifold patterns, namely states, in response to various task demands, enabling cognitive and behavioral flexibility as well as optimal performance (1–6). Alterations in dynamic reconfiguration of functional brain networks when task demands change have been linked to chronic stress-related psychiatric symptoms such as anxiety and depression (7,8), implying the adverse effects of long-term stress on emotional and higher-order cognitive functions (9,10). On the other hand, stress has been considered an online adaptive system that can shape the brain and cognition to assess and deal with the environment (11). Research in animal models also argues that the long-term or chronic stress response represents an adaptive defense mechanism that attempts to cope with the external environment (12). Thus, there is a need to elucidate whether and how long-term stress shapes dynamic reconfiguration of large-scale functional brain networks across multiple task demands involved in emotion and cognition, which can deepen our understanding of stress-related adaptative and maladaptive effects on the human brain.

It has been recognized that functional brain networks are rapidly reconfigured in response to acute stress, characterized by hyper-activation in the emotional, salience, and sensorimotor networks and hypo-activation in the executive control networks (ECN) and default mode networks (DMN) (13–16). In contrary to unpredictable acute stressors, sustained or long-term stressors are often predictable (17), with unique features including prolonged exposure to the same stressor and lasting release or exhaustion of cortisol, a stable indicator of stress-induced neuroendocrine response (18). It thus remains open whether and how such acute stress-induced reconfiguration of brain networks can apply to individuals experiencing long-term stress. Empirical evidence from separate neuroimaging studies converges that long-term stress alters DMN and ECN involved in emotional and cognitive conditions, although there is less agreement on the direction of these alterations. Maladaptive direction often refers to stress-induced impairment and inhibition, such as long-term stress can lead to hypo-activation in the dorsolateral prefrontal cortex (dlPFC), a key node of ECN, during attention control task, which is associated with low performance (19), and reduced connectivity of dlPFC with the amygdala when regulating emotions, which is associated with less capable of down-regulating negative emotion (20). On the other hand, adaptive direction often refers to stress-related benefits and facilitation, such as increased activation in the ventromedial prefrontal cortex (a key node of DMN) under sustained stress implies adaptive coping behavior (21), and greater functional connectivity between the dlPFC and hippocampus (a key node of DMN) under sustained stress make people feel less stressed (22). Research in animal models have also observed chronic stress increases the activity in the prefrontal cortex and hippocampus, which can improve resilience, mood, and cognitive function (23,24). As a result, long-term stress has a profound impact on the brain, emotion and cognition, with either adaptative or maladaptive outcomes seeming different from the reorganization of brain networks under acute stress. Although the effects of long-term stress have been extensively studied from the perspective of brain localization, this approach provides useful information on specific brain regions. The human brain, however, is recognized to work as an open, complex system that enables the nuanced cognitive and affective function, most likely through dynamic interactions among large-scale functional brain networks (25–27). Thus, studying the dynamic configuration of large-scale brain networks can provide valuable insights into the effects of long-term stress on the human brain, and can offer a more comprehensive understanding of the neural mechanisms underlying adaptation and maladaptation.

A fundamental question in the neurobiology of long-term stress is to understand whether and how functional brain networks are adapted to support emotional and cognitive task demands. Most previous studies in long-term stress using single-task paradigms provide limited information about the dynamic reconfiguration of large-scale brain networks across multiple task demands. It is recognized that adapting to varying task demands necessitates the engagement of cognitive control (28), whereby the brain actively coordinates specific neural resources based on current task demands to achieve goal-directed behavior. For example, taking a rest state and performing emotional and working memory tasks requires different organization patterns of large-scale functional brain networks due to their goals differ (29–33). Neuroimaging research has established that the prefrontal cortex plays a crucial role in cognitive control (28), and the frontal-parietal network (FPN) is a flexible hub of cognitive control (34,35). However, long-term stress can shift cognitive control to habitual control with reduced flexibility of learning (36), which may induce maladaptive effects in meeting ever-changing external demands. As mentioned above, previous studies have shown that long-term stress may promote adaptive response, likely through functional interactions between DMN and ECN (37). Notably, the FPN consisting of the lateral parietal cortex and the dlPFC, is part of ECN (34,35). Thus, one may conjecture that individuals exposed to long-term stress may adapt to multiple task demands by allocating more neural resources to the ECN. Although the DMN is not typically associated with cognitive control processes, recent studies have demonstrated that increased activation of the DMN can have positive effects on stress response (22), suggesting a possible stress-buffering effect. This stress-buffering effect may enable individuals to meet the current task demands more effectively and adaptively (23,24). Furthermore, other studies also suggest that increased DMN activation may support adaptive task performance by facilitating the integration of internal and external information (38). However, it still remains unexplored whether and how long-term stress affects the dynamic reconfiguration of DMN and ECN across multiple task demands.

The mainstay of functional neuroimaging studies in long-term stress utilizes conventional analytic approaches to evaluate stress-related changes in activation and connectivity of brain regions and/or networks, by averaging fMRI signals obtained from several-minute scanning. These approaches are sensitive to detect stationary changes in brain activity and connectivity, but offer insufficient information to capture the inherently dynamic nature of functional brain networks (39). Recent evidence shows that latent brain states, reflecting temporal dynamic reconfiguration of certain functional brain networks are critical to detect a variety of emotional and cognitive as well as the resting phase’s brain state dynamics (40–42). In particular, the Hidden Markov Model (HMM) approach, inferring the brain state of each timepoint through the variational Bayes inference (see Methods) (43), is widely used to investigate dynamic brain states at rest and in tasks (44). Recently, a study effectively captured rich brain state dynamics during movie viewing compared to the resting state by using the HMM to estimate the temporally concatenated timeseries (45). Thus, we assume that the HMM can enable us to detect dynamic reconfiguration of functional brain networks across emotional and cognitive task demands together and test the adaptive and maladaptive effect of long-term stress.

Here we aim to investigate how long-term stress affects the dynamic reconfiguration of large-scale functional brain networks across resting state, emotional and working-memory processing, which have different task goals and are conventionally characterized by the involvement of the DMN (31), salience network (SN) (32,46), and ECN (37,47,48), respectively. To this end, we recruited participants who were preparing for China’s National Postgraduate Entrance Examination (CNPEE) as the long-term stress group. This examination is competitive, with a less than 33% acceptance rate, and college students have to exert effort for at least six months to pursue their master graduate (Duan et al., 2013; Liu et al., 2021), which has been proven effective to induce chronic psychological stress (19,51). As the control group, we recruited college students who matched in age and sex and did not prepare for the CNPEE or suffered any stressful events. To account for the multifaceted nature of long-term stress, such as social, psychological, and physiological/physical responses (7,8,52–54), participants were asked to measure negative emotions, bodily and organic symptoms, as well as interpersonal relationships on day 1 (**Fig. 1A**). On day 2, participants underwent an fMRI study while taking a resting state and performing emotional matching task and working memory in sequence (**Fig. 1A**), and the HMM was used to determine the dynamic configuration of functional brain networks during these different demands under long-term stress and controls. Based on the aforementioned empirical observations of stress-related effects on emotion and cognition, we expected long-term stress would shape the dynamic configuration of DMN and ECN to flexibly and adaptively meet multiple task demands.

**Fig. 1.**
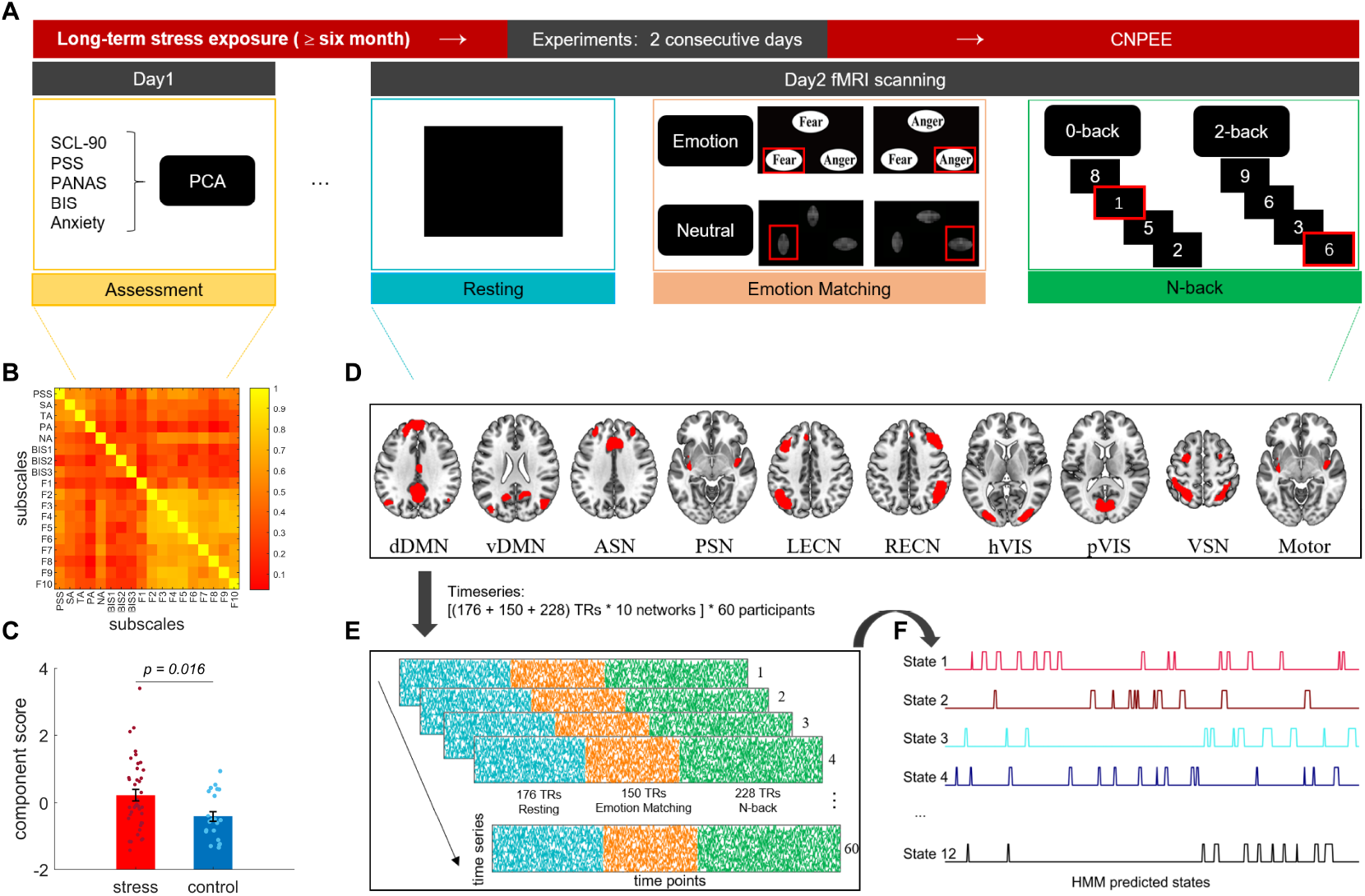
Experiment design and a schematic illustration of HMM analysis. **A**. Experiment design. Participants who suffered long-term stress (at least 6 months) were recruited 1-3 weeks before the end of the major exam stressor, and who did not suffer long-term stress were recruited as the control group. All participants completed five psychological assessments consisting of 18 subscales (Day 1) and took a resting state and performed emotion matching task and working memory (N-back) in sequence in fMRI scanning (Day 2, see Methods). **B**. Matrix graph depicts the correlation coefficients of 18 subscales of five psychological assessments, and most of them showed high correlations, *P*_FDR_ < 0.05. **C**. The bar graph shows a significant difference in long-term stress (the first component score extracted from 18 subscales, Y axis) between the stress and control groups. **D**. Schematic illustration of ten brain sub-networks that were considered regions of interest. **E**. Timeseries (Y axis) were extracted from them (X axis, a total of 554 time points: 176 TRs for resting, 150 TR for emotion matching, and 228 TR for N-back) of all 60 participants during the stress (*N* = 39) and control (*N* = 21) groups. **F**. 12 brain states were inferred through the Hidden Markov Model (HMM), and each time point was categorized into a single brain state. SCL-90, Symptom Checklist-90. PSS, perceived stress scale. PANAS, positive affect, and negative affect scale. BIS, Barratt Impulsiveness Scale. Anxiety, State-Trait Anxiety Inventory (STAI). SA, state anxiety. TA, trait anxiety. PA, positive affect. NA, negative affect. F1-F10, subscales in SCL-90. dDMN and vDMN, dorsal and ventral Default Mode Networks. ASN and PSN, Anterior and Posterior Salience Network. LECN and RECN, Left and Right Executive Control Networks. pVIS, Primary Visual Network. hVIS, High Visual Network. VSN, Visuospatial Network. Motor, Sensorimotor Network.

## Results

### Long-term stress exposure leads to psychological distress

First, we examined psychological measurements for participants who were exposed to long-term exam stress as compared to controls. We implemented principal component analysis (PCA) for five assessments consisting of 18 self-reported psychological, physical, and social subscales that are closely associated with stress to identify the major representative component (see Methods). Most subscales are positively correlated with each other (all *p* < 0.05, **Fig. 1B**), indicating they tend to assess a similar construct of psychosocial stress. This analysis revealed a first principal component that explained 48% of the variance, with positive loadings in all subscales except the positive affect subscale in PANAS (−0.35) (Supplementary **Fig. S1**). This indicates that participants with high scores of the first principal component exhibited lesser values in the positive affect subscale. Hence, the first principal component is assumed to characterize the negative psychological distress in response to long-term exam stress in our present study. We examined the difference between the stress and control groups in the first component scores. As expected, an independent sample t-test revealed that participants in the stress group (0.22 ± 1.07) exhibited significantly greater scores of long-term stress as compared to those in the control group (−0.42 ± 0.71) (*t*_(58)_ = 2.47, *P* = 0.016, Cohen’s d = 0.70, 95% CI = 0.12, 1.16) (**Fig. 1C**). These results indicate the effective induction of long-term stress in participants who were exposing the CNPEE preparation.

### Distinct latent brain states are dominant to rest, emotional, and cognitive tasks

In the following analysis, we aimed to identify the configuration of functional brain networks during the resting state, emotional and WM processing by collapsing across stress and control groups. To achieve this, we applied the HMM on the temporally concatenated timeseries of resting state, emotional and WM tasks extracted from ten brain sub-networks of interest (55) (**Fig. 1D-E**). The ten sub-networks considered in the present study belong to the DMN, ECN, SN, and sensory network, which have been shown to play a crucial role at rest and in emotional and cognitive tasks. This HMM analysis yielded twelve brain states (**Fig. 2A**) that were inferred by the variational Bayes inference. Each time point was categorized into a single brain state (**Fig. 1F**) characterized by distinct activation intensity of ten sub-networks, with three key metrics: 1) fractional occupancy, the total proportion of time spent in a brain state; 2) mean lifetime, the average time spent in a brain state before switching to the next brain state; 3) transition probability, the probability of switching from one brain state to another one.

**Fig. 2.**
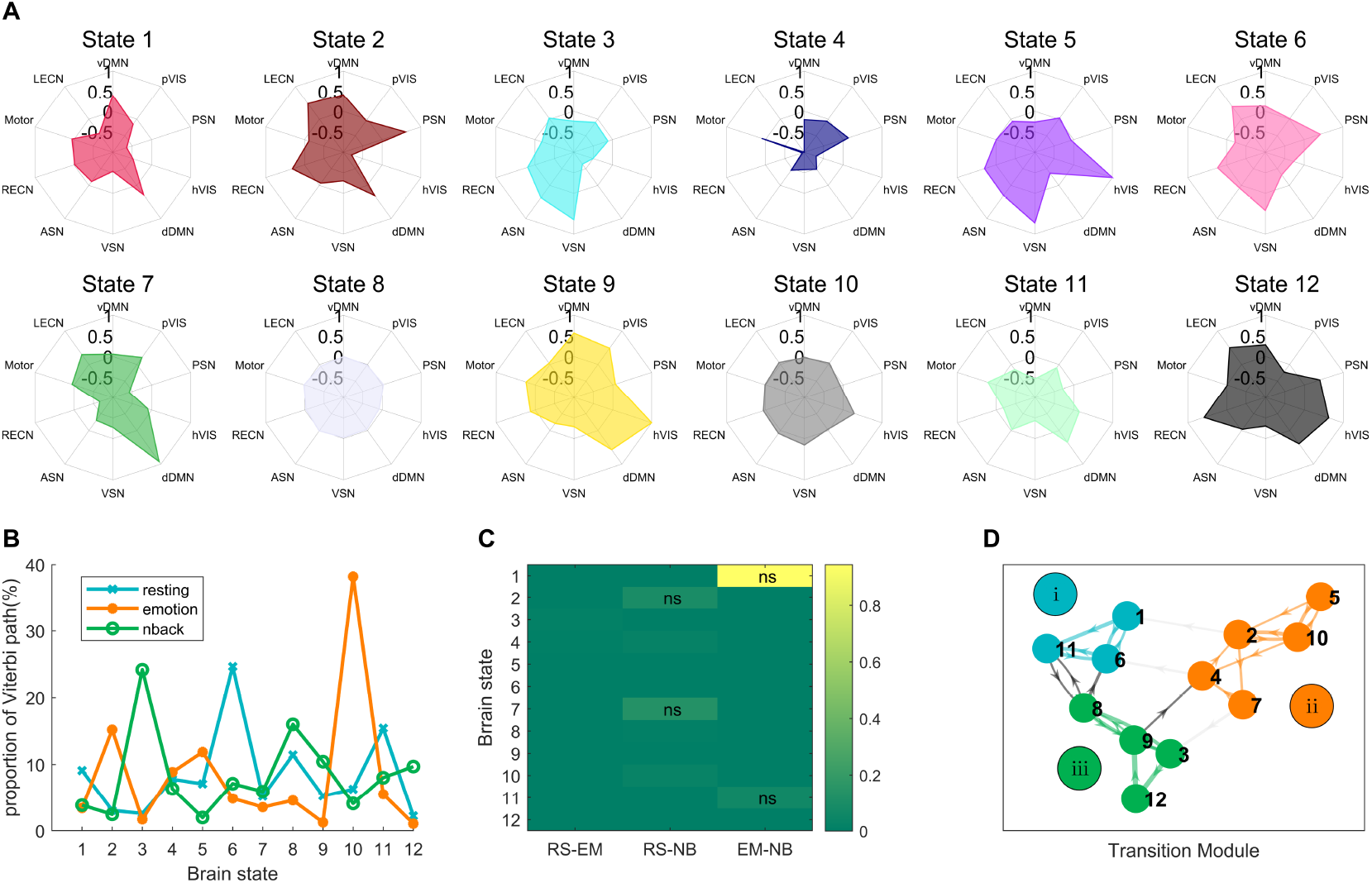
Latent brain states involved in rest, emotional, and working memory processing. **A**. Polar plots depict the 12 brain states with different activation of ten sub-networks as mentioned in Fig.1. **B**. The total proportion of occurrence for each brain state in each task. The Viterbi path is the most likely sequence of states across time points. **C**. Almost all brain states have a significant difference in fractional occupancy between any two tasks. RS, resting. EM, emotion. NB, N-back. Each rectangular indicates an FDR-corrected p-value, and ‘ns’ indicate the FDR-corrected p-values are greater than 0.05. **D**. Three modules were obtained by applying the Louvain community detection algorithm on the top 21% transition at the group level. Arrow thickness shows the probability of a transition from one state to another. Roman numerals indicate module 1, module 2, and module 3, respectively.

We then tested the differences in the three key metrics of brain states across different task demands by collapsing across the stress and control groups. As shown in **Fig. 2B**, we found State 6 as the dominant brain state (25%) at rest, characterized by relatively high BOLD activity in subnetworks of the vDMN, ECN, PSN, and VSN; State 10 as the dominant brain state (38%) in the emotion matching task, with high activity in visual subnetworks of the hVIS and pVIS; State 3 as the dominant brain state (24%) in WM task, with high activity in subnetworks of the RECN, ASN, and VSN. Our further analyses confirmed significant differences in fractional occupancy between rest, emotion, and WM tasks in almost all brain states (*P*_FDR_ < 0.05, **Fig. 2C**). Subsequently, we applied the Louvain community detection algorithm (56) to the top 21% transition probability matrix acquired from all participants and all tasks. We found three modules (**Fig. 2D**) corresponding rest, emotion and WM tasks, respectively. In particular, we calculated the total transition probability from other states except for self-switching for each brain state and used them as features for support vector machines classification of tasks (see Methods). As expected, the accuracy based on module 1, named Module_rest_, is significantly higher than the other modules when classifying the resting state in all three tasks (Supplementary **Fig. S2**). Likewise, module 2 and module 3, named Module_emotion_ and Module_WM_, performed best when classifying the emotion and WM tasks, respectively. It is worth noting that the dominant states at rest and in emotion and WM tasks belong to Module_rest_, Module_emotion_, and Module_WM_. It is thus reasonable to assume these modules as task-specific modules and states in each module as task-specific states, named State_rest_ (consisting of states 1, 6, and 11, which exhibit shared activation of the DMN), State_emotion_ (consisting of states 2, 4, 5, 7, and 10, which exhibit shared activation of the SN, except for State 7), and State_WM_ (consisting of states 3, 8, 9, and 12, which exhibit shared activation of the ECN, except for State 8) respectively. These results indicate that the configuration of brain networks is task-specific in this study, highlighting the brain’s ability to adapt to changing task goals and neural demands. This unique configuration of brain networks for each task demand allows us to estimate how long-term stress impacts this adaptation process.

### Long-term stress prolongs brain states linked to task demands and cognitive control

To explore the effects of long-term stress on the dynamic reconfiguration of functional brain networks across multiple task demands, we conducted a comprehensive analysis of the resting state, emotion, and working memory tasks. Separate independent sample t-tests were conducted for dynamic properties of brain states at rest, emotion and WM tasks between the long-term stress and control groups. These analyses revealed higher fractional occupancy (*t*_(58)_ = 3.14, *P* = 0.003, Cohen’s d = 1.13, 95% CI = 0.13, 0.57, *P*_FDR_ < 0.05) and longer mean lifetime (*t*_(58)_ = 2.84, *P* = 0.006, Cohen’s d = 0.72, 95% CI = 0.23, 1.31, *P*_FDR_ = 0.072) of state 1, named State_DMN_, at rest in the stress group than controls (**Fig. 3A-B**). Notably, this state belongs to State_rest_ and is characterized by high dDMN and vDMN (**Fig. 2A**), which are well-defined as resting-like activity. These results indicate that long-term stress prolongs the brain state related to task demands at rest.

**Fig. 3.**
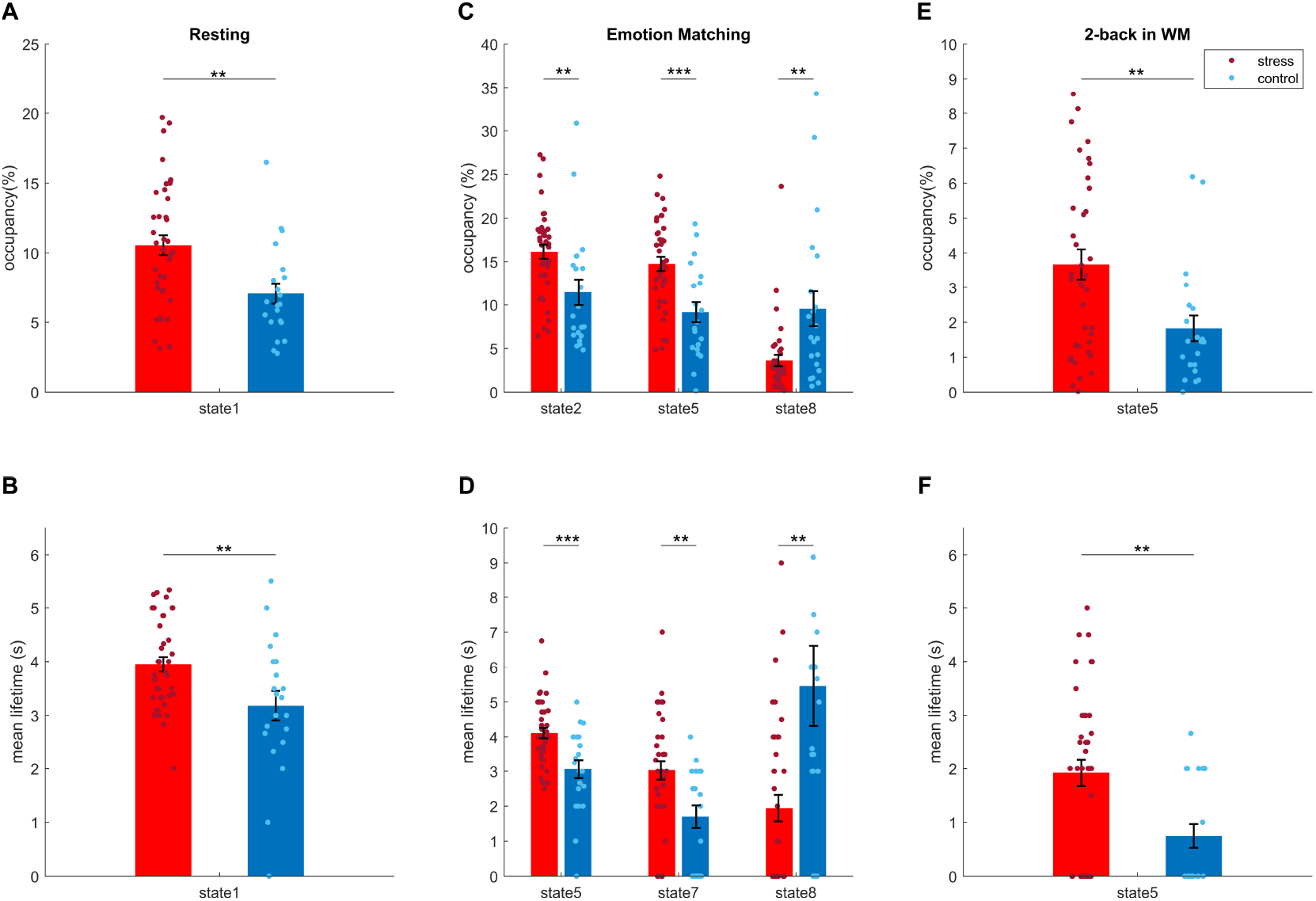
Effects of long-term stress on latent dynamic brain states during rest, emotional and working-memory processing. The top panel corresponds to the fractional occupancy in each task, whereas the bottom panel corresponds to the mean lifetime in each task. **A-B**, Resting. State 1 (State_rest_) shows higher occupancy and mean lifetime in the stress group than controls. **C-D**, Emotion matching. States 2, 5 (State_emotion_) have higher occupancy in the stress group, whereas state 8 has higher occupancy in the control group. The mean lifetime of states 5 and 7 (State_emotion_) is higher in the stress group, while the mean lifetime of state 8 is higher in the control group. **E-F**, 2-back. State 5 (State_emotion_) shows higher occupancy and mean lifetime in the stress group compared to controls. **, *p* < 0.01; ***, *p* < 0.001.

During emotion matching task, we found that long-term stress prolongs brain states related to emotion processing and cognitive control. In particular, participants under long-term stress showed higher fractional occupancy in state 2 (*t*_(58)_ = 3.04, *P* = 0.004, Cohen’s d = 0.82, 95% CI = 0.02, 0.08, *P*_FDR_ < 0.05) and state 5 (*t*_(58)_ = 4.04, *P* < 0.001, Cohen’s d = 1.20, 95% CI = 0.03, 0.08, *P*_FDR_ < 0.05) (**Fig. 3C-D**). Moreover, the mean lifetime of state 5 (*t*_(58)_ = 3.69, *P* < 0.001, Cohen’s d = 0.96, 95% CI = 0.48, 1.61, *P*_FDR_ < 0.05) and state 7 (*t*_(58)_ = 3.03, *P* = 0.004, Cohen’s d = 0.84, 95% CI = 0.45, 2.21, *P*_FDR_ < 0.05) in the stress group is significantly larger than controls (Fig. 4d). It is worth noting that the fMRI signals defining states 2, 5, and 7 that belongs to State_emotion_ loads on functional networks supporting emotion processing (SN and Visual networks) and cognitive control (ECN) (**Fig. 2A**). Furthermore, the DMN also exhibits high activation in states and 7, which may play a role in regulating negative emotions induced by negative stimuli (e.g., Goldfarb et al., 2020). In contrast, the stress group’s fractional occupancy (*t*_(58)_ = -2.82, *P* = 0.009, Cohen’s d = -0.86, 95% CI = -0.09, -0.03, *P*_FDR_ < 0.05) and mean lifetime of state 8 (*t*_(58)_ = -3.58, *P* = 0.001, Cohen’s d = -0.86, 95% CI = -5.49, -1.55, *P*_FDR_ < 0.05) is significantly smaller than controls (**Fig. 3C-D**). State 8 exhibits relatively uniform negative fMRI signals across ten sub-networks (**Fig. 2A**), implying no activation of specific function networks. Interestingly, the pattern of the above results also occurred during separate emotional and neural conditions in emotion matching task (Supplementary **Fig. S3**). We further implemented network-based prediction analysis (see Methods) to estimate the difference in transition probability in the whole emotion matching task between long-term stress and control groups. Compared to controls, the stress group showed a significantly higher probability of transition from states 8 and 10 to states 2, 3, 5, 7, and 12 than the control group (Supplementary **Fig. S4**, *P*_FWE_ < 0.05). Participants in the control group, however, exhibited more likely to transit from states 2, 3, and 5 to state 8 (Supplementary **Fig. S4**, *P*_FWE_ < 0.05). These results again indicate that the stress group spent more time in states 2, 5, and 7 from State_emotion_ with high activation of SN, DMN, ECN, and visual networks, which can support people in processing emotions and cognitive control in adapting to the new emotional task from the former resting phase.

**Fig. 4.**
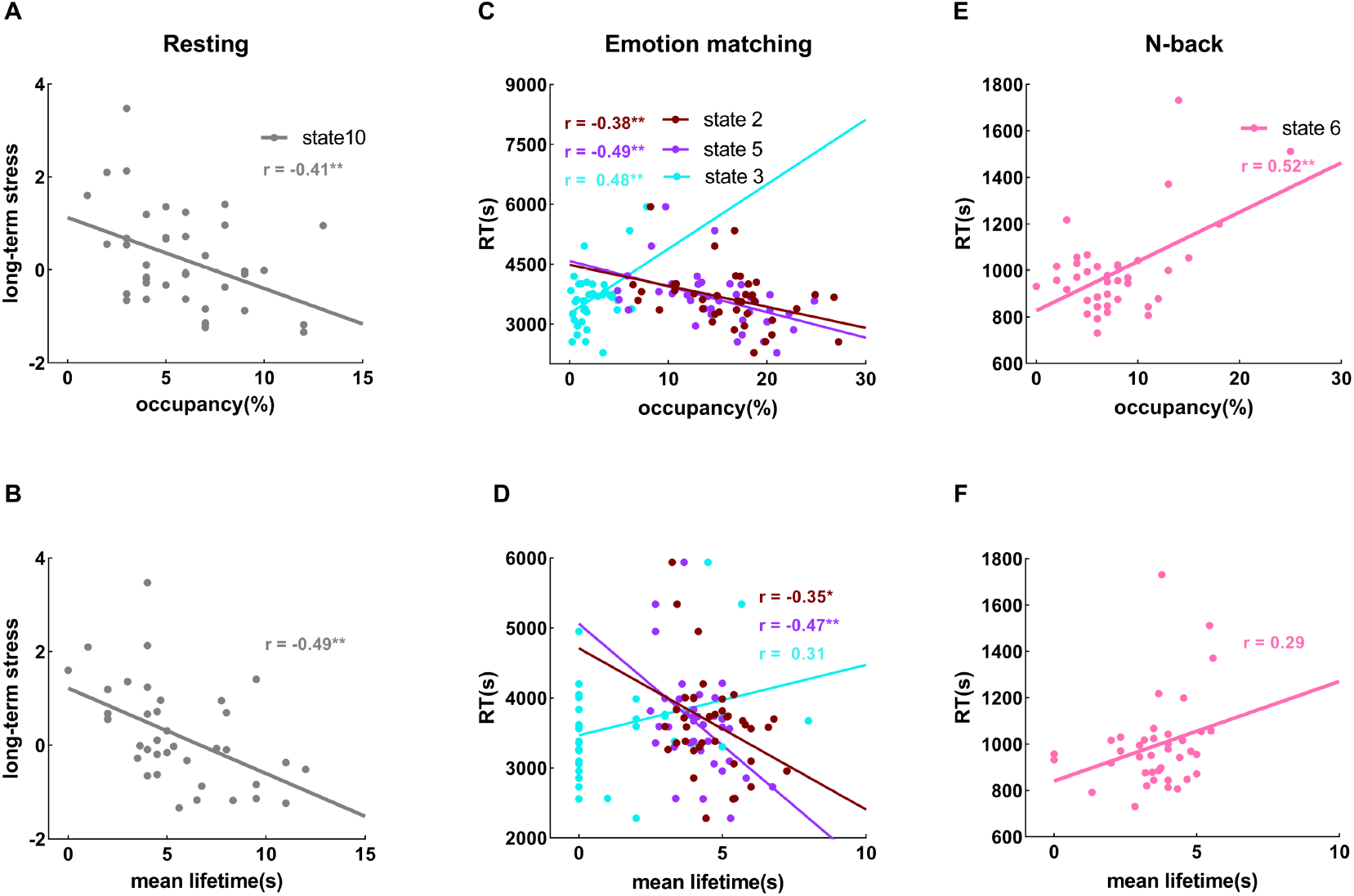
The relation between brain state dynamics and psychological distress and behavioral performance under long-term stress. The top panel corresponds to the fractional occupancy in each task, whereas the bottom panel corresponds to the mean lifetime in each task. **A-B**, Resting. Long-term stress negatively correlates with the fractional occupancy and mean lifetime of state 10 (State_emotion_). **C-D**, Emotion. The occupancy and mean lifetime of state 2 and state 5 (State_emotion_) are negatively correlated with the reaction time, whereas the occupancy and mean lifetime of state 3 (State_WM_) are positively correlated with it. **E-F**, N-back. The reaction time is (marginally) positively correlated with the occupancy and mean lifetime of state 6 (State_rest_). *, *p* < 0.05; **, *p* < 0.01.

Similar to the emotion matching task, we observed higher fractional occupancy (*t*_(58)_ = 2.84, *P* = 0.006, Cohen’s d = 0.78, 95% CI = 0.01, 0.03, *P*_FDR_ = 0.072) and longer mean lifetime (*t*_(58)_ =3.10, *P* = 0.003, Cohen’s d = 0.89, 95% CI = 0.42, 1.94, *P*_FDR_ < 0.05) of State 5 in the stress group than controls (**Fig. 3E-F**) during the 2-back condition. This state is characterized by high activation of visual networks, RECN, and ASN, which may support stressed people to process digits stimuli and perform working memory at a high cognitive load with high cognitive control. However, state 5 is not from State_WM_, which may indicate that individuals under long-term stress stick to the neural response of the former emotion matching task. On the other hand, state 5 may be a more fundamental brain state supporting emotional and cognitive tasks (see Discussion).

In sum, these results indicate that long-term stress shapes the dynamic reconfiguration of large-scale brain networks dominant to different task demands and cognitive control, which differs from previous observations on acute stress that deactivates DMN and ECN, signaling the unique neural adaptation under long-term stress.

### Brain state dynamics are linked to psychological distress and behavioral performance under long-term stress

We further tested the relationships between brain state dynamics and psychological measures of long-term stress as well as task performance in the stress and control groups. At rest, we found a significant negative correlation between the mean lifetime of State10 named State _visual_ with high activation of visual networks and psychological distress under long-term stress (*r*_(39)_ = -0.49, *P* = 0.002, *P*_FDR_ < 0.05) (**Fig. 4B**). The fractional occupancy of State _visual_ showed a similar effect (*r*_(39)_ = -0.41, *p* = 0.009), though it reaches no significant after FDR correction (*P*_FDR_ = 0.11) (**Fig. 4A**). Moreover, permutation testing (see Methods) was used to assess the null hypothesis of equality in these correlations between long-term stress and control groups. We found that the mean lifetime (*p* < 0.001) of State _visual_ at rest in the stress group was significantly more correlated with psychological distress than controls. Likewise, we found a marginally significant trend for higher fractional occupancy in stress than controls (*p* = 0.077) (Supplementary **Fig. S5**). In particular, State _visual_ at rest is not of State_rest_ (see **Fig. 2D**), and so these results indicate people who spent more time in the non-task-specific state showed less psychological distress. We observed no reliable correlation between other brain states at emotional and working memory tasks with psychological distress in the stress or control groups.

Interestingly, we observed that the stress group’s fractional occupancy and mean lifetime of State_emotion_, such as State 2 (*r*_(39)_ = -0.375, *P*_uncorr_ = 0.019; *r*_(39)_ = -0.352, *P*_uncorr_ = 0.028) and State 5 (*r*_(39)_ = -0.458, *P*_uncorr_ = 0.003; *r*_(39)_ = -0.466, *P*_uncorr_ = 0.003) are negatively correlated with RTs, while fractional occupancy and mean lifetime of State_WM_, such as State 3 showed a positive correlation with RTs (*r*_(39)_ = 0.437, *P*_uncorr_ = 0.005); *r*_(39)_ = 0.307, *P*_uncorr_ = 0.058) (**Fig. 4C-D**) in the emotion matching task. During the WM task, the fractional occupancy and mean lifetime of State_rest_, such as State 6 is (marginally) positively correlated with the RT (*r*_(39)_ = 0.522, *P*_uncorr_ = 0.001; *r*_(39)_ = 0.290, *P*_uncorr_ = 0.073) (**Fig. 4E-F**). These results may support that people under long-term stress perform tasks better when spending more time in task-specific brain states while worst when spending more time in non-task-specific brain states. We further tested the difference in correlation effects between long-term stress and control groups. Permutation testing again revealed significantly more correlations of fractional occupancy and mean lifetime with behavioral performance in the stress group compared with controls (Supplementary **Fig. S5**). However, there is no significant difference in correlations between the two groups in the dynamics of State 5.

Together, prolonged (non) task-specific brain states are (positively) negatively related to task performances, whereas prolonged non-task-specific brain state is negatively related to psychological distress, with better prediction accuracy in the stress group than in controls. These results indicate that exposure to long-term stress leads to increased sensitivity of task-driven brain states, which implies both adaptive and maladaptive changes on psychological and neurobehavioral levels.

## Discussion

In this study, we investigated the effect of long-term stress on dynamic reconfiguration of large-scale functional brain networks across multiple tasks. Long-term exam stress resulted in higher psychological distress compared to controls. Critically, individuals experiencing long-term stress spent more time engaging DMN-centric brain states during the resting state and more time engaging visual-, SN-, ECN-, or DMN-centric brain states during subsequent emotional and working memory tasks. Interestingly, brain state dynamics at rest were linked to psychological distress during the stress group, while brain state dynamics in emotional and cognitive tasks are highly correlated with behavioral performance, with the correlation direction depending on the brain states whether are of task-specific states or not. Furthermore, long-term stress showed higher correlation effects than the control group, implying increased sensitivity to specific brain networks. Together, our findings suggest that long-term stress shapes brain networks’ reconfigurations in response to different task demands or goal-directed behavior and cognitive control, in ways that differ from those observed under acute stress. This provides essential implications for neurocognitive flexibility and (mal)adaptation under long-term stress.

### Functional organization of brain networks varies based on task demands

In line with previous findings of the flexible reconfiguration of brain networks when adapting to varying task demands (1–5), the dominant latent brain states at rest and during tasks in this study differ, indicating our successful manipulation of different task demands. In particular, State_rest_ consisting of states 1, 6, and 11 exhibits shared activation in the DMN, a crucial brain network supporting the resting state (31). In contrast, State_emotion_, which consists of states 2, 4, 5, 7, and 10, shows shared activation in the SN, except for state 7, which is a key brain network for processing emotions (32,46). During working memory task, State_WM_ consisting of states 3, 8, 9, and 12 exhibits shared activation in the ECN, except for state 8, which is a crucial brain network for dealing with cognitive loads (37,47,48). In addition, extensive evidence suggests that the FPN, a part of ECN, is a hub of cognitive control in adapting various tasks (34,35). Thus, the ECN not only serves a specific function for current task demands but also serves as general cognitive control in switching to new tasks from the former conditions. As discussed below, long-term stress shapes the dynamic reconfiguration of brain networks relevant to multiple task demands and cognitive control.

### Long-term stress increases the persistent time and transitions to brain states relevant to task demands and cognitive control

One of our major results is that long-term stress has significant effects on the reconfiguration of brain networks relevant to task demands and cognitive control. At rest, individuals under long-term stress spent more time in the DMN-centric brain state (state 1), which is in line with previous findings (51). The DMN is known to activate during the resting state when individuals focus on their internal state (31). As mentioned above, State 1 is of State_rest_ (**Fig. 2D**), which can support individuals in fulfilling the requirements of the resting phase in the scanner, such as staying awake and not thinking about anything in particular. Thus, the hyperactivation of DMN may suggest stressed individuals allocate more task-specific neural resources to meet current task demands, which may represent a possible adaptive neural response to long-term stress.

As for the emotion matching task, individuals experiencing long-term stress spent more time in State_emotion_, such as states 2, 5, and 7, which show high loadings of fMRI signals on functional networks supporting emotion processing (SN and visual network) and cognitive control (ECN). The increased activation of SN and visual network in stressed individuals could potentially be a neural adaptation to meet the task demand of processing emotions (32,46). Similarly, increased activation of the ECN may be a neural adaptation to improve the ability of cognitive control and further promote the transition from the resting phase to the emotion matching task (34,35). Furthermore, the DMN also exhibits high activation in states 2 and 7 and the core brain regions of DMN and ECN have been shown to play an essential role in regulating emotions (22,57–59) and their functional interactions may promote adaptive response under long-term stress (37). Spending more time in states 2, 5, and 7 may thus help stressed individuals regulate the higher activation of SN induced by long-term stress and negative emotional stimulus, enabling them to achieve similar task performance to controls (Supplementary **Fig. S6**). Our study further found that stressed individuals more often switched to states 2, 5, and 7 from other brain states. This suggests that they may shift the dynamic configuration of function networks relevant to task demands and cognitive control, adapting to the emotion matching task from the resting state. Finally, our study observed similar effects in both emotional and neutral conditions (Supplementary **Fig. S3**), consistent with the fact that daily stress and emotions evoked in the task will carry over subsequent neutral blocks and affect perception (60), decision (61,62), and memory (63).

Likewise, our results further indicate that stressed people spend more time in state 5 during the higher-load 2-back condition. Although state 5 is characterized by high activation in the visual network, right executive control networks (RECN), and anterior salience networks (ASN), which are crucial functional brain networks supporting digit processing and working memory, state 5 is not part of State_WM_. One possible explanation for this observation is that state 5 may represent a fundamental brain state that supports both emotional and cognitive tasks. Alternatively, it is plausible that individuals under long-term stress may rely on the habitual neural response of the previous emotion matching task when performing the working memory task, a phenomenon observed in previous studies showing that long-term stress promotes habitual control (36). However, this temporary habitual control may be adaptive in a performance like under acute stress (64,65), since the prolonged state 5 can support working memory as mentioned above.

### Long-term stress increases sensitivity to the dynamics of task-driven brain states

Another important result revealed that fractional occupancy and mean lifetime of a specific brain state at rest are correlated with psychological distress under long-term stress. This finding is consistent with previous studies showing that the fractional occupancy of brain states at rest can predict overall mental health (66) and mental disorders (42). Specifically, we found that fractional occupancy and mean lifetime of state 10 are negatively correlated with psychological distress under long-term stress, which is characterized by high activation of Visual networks, ECN, and ASN. Although it is believed that intrinsic spontaneous neural activity of DMN reflects the idiosyncratic self, consisting of long-term memories, beliefs, and emotions (67), we found that state 10, which does not involve activation of the DMN, is linked to psychological distress under long-term stress. This may reflect that stressed individuals are more prone to hyper-activation of SN and ECN: the ASN, with its core region in the amygdala, is involved in processing emotional information, while the ECN, with its core region in dlPFC, is involved in cognitive control (68). Moreover, the hyper-activation of visual networks in state 10 is somewhat consistent with previous studies: in rodents, severe stress can induce brain atrophy in the visual cortex (69), and the visual cortex also showed weaker spontaneous activity at rest in PTSD patients (70). Hence, individuals experiencing long-term stress may be more likely to suffer less distress with higher activation of visual networks at rest, which is proved by our results that the hyper-activation of visual networks in state 10 at rest is negatively correlated with long-term stress. Our findings also suggest that there are significant differences in this correlation effect of state 10 between stress and control groups, implying stressed individuals are more sensitive to the dynamic configuration of specific functional brain networks.

Expanding previous findings in emotional and working memory processing (40,41), our results suggest that brain state dynamics in emotional and working memory tasks are more related to behavioral performances during the stress group. Specifically, we found that in the presence of long-term stress, fractional occupancy of two brain states belonging to State_emotion_ is negatively correlated with RTs, whereas fractional occupancy of a brain state belonging to State_WM_ is positively correlated with RTs during emotion matching task. In addition, we observed a positive correlation between the fractional occupancy of a brain state belonging to State_rest_ and RTs during WM task under long-term stress. Thus, spending more time on task-specific brain states (e.g., State_emotion_) may lead stressed people to perform task more quickly, while spending more time in non-task-specific brain states may lead to longer task completion times. Again, permutation testing indicates that these effects are stronger in the long-term stress group compared to controls, implying that stressed individuals are more sensitive to the dynamics of task-specific and non-task-specific brain states, which may be both adaptive and maladaptive neural response under long-term stress.

Together, our findings expand previous research on long-term stress by highlighting that long-term stress shapes dynamic reconfiguration of large-scale brain networks relevant to task demands and cognitive control in different ways compared to acute stress that deactivates DMN and ECN and also makes individuals more sensitive to task-driven functional brain networks.

There are several limitations of our present study. First, we only recruited male participants to avoid potential confounds in menstrual cycles (47) and gender differences in chronic stress perception (71). It is unknown whether these findings could be generalized into a female population. Second, we tested the hypothesis on the long-term exam stress, and it is an open question whether other long-term stress (e.g., financial pressure) would induce similar effects. Finally, we only estimated several functional brain networks of interest. Future studies can explore the whole brain or specific brain regions.

## Conclusions

Our study elucidates that long-term stress can shape dynamic reconfiguration of large-scale functional brain networks across different tasks based on current demands and cognitive control and increase the sensitivity of brain states to psychological distress and task performance. Our findings provide a neurocognitive framework for better understanding the impact of long-term stress on the human brain, with both adaptive and maladaptive changes on psychological and neurobehavioral levels.

## Acknowledgements

This work was supported by the National Natural Science Foundation of China (32130045, 82021004), the Open Research Fund of the State Key Laboratory of Cognitive Neuroscience and Learning (CNLZD1503).

## Data and code availability

Behavioral and fMRI data as well as the analysis code used in this study will be made available upon publication in a peer-reviewed journal.

## Methods

### Participants

We recruited sixty-one healthy male college students for this study. Forty participants (age range: 20-24 years old, mean ± S.D. = 21.63 ± 0.80) in the long-term stress group were preparing for China’s National Postgraduate Entrance Examination (CNPEE). They came to the laboratory for two days, 1-3 weeks before the exam. More information about the screening criteria for long-term exam stress can be found in our previous study (37). Twenty-one participants (age range: 21-23 years old, mean ± S.D. = 21.67 ± 0.71) who did not suffer long-term stress were recruited as the control group. One participant in the stress group was excluded because of failure to finish all tasks during the fMRI scanner, and the left participants were included based on the mean framewise displacement (FD) < 0.22 mm (only one participant’s mean framewise displacement in the WM task is more than 0.2 mm) with proportion of spikes (FD > .25 mm) < 20%, and no spikes above 5 mm were observed (33,72,73). It is worth noting that there is no significant group difference in the mean FD and results in this study keep unchanged after regressing out the mean FD (Supplementary **Fig. S7**).

### Experimental design and procedure

The experiment comprised two days. On the first day, participants completed several self-report psychological assessments: the Perceived Stress Scale (PSS) (74) measuring the participants’ stress level in the last month, the Symptom Checklist-90 (SCL-90) (75) measuring the participants’ psychological and physical health degree in the last week, the Barratt Impulsiveness Scale (BIS) (76) measuring the participants’ impulsiveness in three different dimensions, the State-Trait Anxiety Inventory (STAI) (77) measuring the participants’ state and trait anxiety, and the Positive and Negative Affect Schedule (PANAS) (78) measuring the participants’ subjective mood. On the second day, participants were asked to do three tasks inside the fMRI scanner, including resting, emotion matching, and N-back, and they firstly practiced the last two tasks outside the scanner.

During the resting phase, participants were instructed to relax, remain still with eyes open, state wake up, and not think about anything in particular during the 6-minute scan. During the emotion matching task, we displayed participants with three emotion faces (emotion condition) or three shapes (neutral condition) in each trial. They were asked to select one of two stimuli in the bottom row, which should be in the same category as the target stimuli in the top row. The task consisted of 5 emotion and 5 control blocks, and there was a total of 6 trials (5s) in each block. During the N-back task, we utilized 0-back and 2-back conditions indicated by a 2-s cue, with 6 blocks in each condition. Participants saw 15 digits in a pseudorandomized sequence in each block, and one digit was presented for 400ms, followed by an inter-stimulus interval of 1400ms. Participants were asked to detect whether the current digit was ‘1’ in the 0-back condition and whether the current digit had appeared two positions back in the sequence in the 2-back condition. Participants were told to press the button when the target occurred as fast as possible during each task.

### Long-term stress

We computed a total of 18 subscales from all of the assessments: one in PSS, which was the sum of items; ten in SCL-90, which were the quotient between the sum of items and the number of items in each subscale; three in BIS, which were the sum of items in each subscale; two in STAI, which were the sum of items in each subscale; and two in PANAS, which were the mean of items in each subscale. For these subscales, we replaced the missing values with the mean value of the sixty participants. The Principal Component Analysis (PCA) was realized using Dimension Reduce in IBM SPSS Statistics version 22.0. We used the PCA method and the max equation rotation to achieve the Factor Analysis. To infer the long-term stress, we utilized the individual scores of the first PCA component to represent the participants’ general psychological distress or long-term stress (see Results).

### Image acquisition and preprocessing

Scanning was performed with a 3T TRIO MRI scanner located at the National Key Laboratory of Cognitive Neuroscience and Learning, Beijing Normal University. Functional brain images were acquired using a gradient-recalled echo-planar imaging (GR-EPI) sequence (33 slices; repetition time, 2000ms; echo time, 30ms; flip angle, 90?; field of view, 200 × 200 mm; voxel size, 3.1 × 3.1 × 4.6 mm). The resting task consisted of 180 volumes (6 min), the emotion matching task consisted of 154 volumes (5 min), and the N-back task consisted of 232 volumes (8 min). High-resolution anatomical images were acquired through a 3D sagittal T1-weighted magnetization-prepared rapid gradient-echo sequence (192 slices; repetition time, 2530ms; echo time, 3.45ms; flip angle, 7?; field of view, 256 × 256 mm; voxel size, 1 × 1 × 1 mm).

Functional brain images were preprocessed using SPM12 (https://www.fil.ion.ucl.ac.uk/spm/software/spm12/) based on MATLAB platform. The first four volumes in each task were discarded to allow for signal equilibrium. The remaining images were corrected for slice acquisition timing, realigned for head motion correction, spatially normalized into the MNI space (version, 2009c), resampled into 2-mm isotropic voxels, and smoothed by convolving a 6-mm Gaussian kernel. Finally, we implemented bandpass filtering (0.01-0.15 Hz) (45)

### Brain Networks and Time series

Time series were extracted from 10 of 14 canonical brain networks defined by cognitive states with whole-brain connectivity patterns (79). These 10 brain networks consist of two subnetworks of DMN, two subnetworks of SN, two subnetworks of ECN, three subnetworks of the visual network, and a sensorimotor network. It is evident that we selected the first six subnetworks because they are highly involved in the current tasks. Moreover, we decided on the last four networks due to emotional and cognitive tasks requiring the engagement of vision and motion brain regions. Therefore, including brain networks ensures the HMM considers important brain activities in this study. For each participant, task, and brain network, mean BOLD time series were extracted and further demeaned and detrended. Then, we implemented regression of 24 motion parameters, global signals in white matter, and cerebrospinal fluid. After normalizing (unit standard deviation), we obtained three matrices for each participant, including 176 time points × 10 brain networks in resting, 150 time points × 10 brain networks in emotion matching, and 228 time points × 10 brain networks in N-back.

### Hidden Markov Model

The time series of the three tasks were firstly temporally concatenated for each participant, and each participant obtained 554 time points × 10 networks. The 60 participants were temporally concatenated secondly, resulting in an fMRI data matrix of 33240 time points × 10 networks. Finally, we inferred brain states by fitting a Hidden Markov Model (HMM) to the fMRI data matrix. The HMM assumes that time series data is produced by a sequence of hidden brain states (43). Each brain state was described as a multivariate Gaussian distribution, which represented the activation pattern of the mean BOLD signal across the 10 brain networks. The HMM mainly generated three parameters to assess the brain state dynamics: fractional occupancy, mean lifetime, and transition probability. Fractional occupancy refers to the proportion of time the HMM spent in a state; mean lifetime refers to the average time the HMM spent before switching to another state; transition probability refers to the probability of switching from one state to another.

We used HMM-MAR toolbox(https://github.com/OHBA-analysis/HMM-MAR) (43) to estimate the HMM parameters and inferred 12 states with an input of 33240 time points × 10 networks data matric. The number of states was decided according to the free energy (80). Previous HMM studies had also confirmed that 6-12 states were appropriate for describing brain function temporal dynamics (43,45,81–83). Beyond the group-level fractional occupancy, mean lifetime, and transition probability, we calculated each of these three parameters for each participant during each task (see HMM-MAR wiki). Therefore, we can examine the difference between the stress and control groups regarding the individual-level measures.

### Task Classification based on transition modules

First, the Louvain community detection algorithm (56) was applied to the top 21% transition matrix acquired from all participants. According to the weights between any two nodes (e.g., the transition probability from one state to another), this algorithm can detect communities where nodes display robust connectivity with a community. We found three communities (modules) through the function of modularity maximization. Second, the support vector machine (SVM) was used to classify tasks based on these three modules. We calculated the total probability of transition from other states to states in modules as classification features, representing the necessity for one state to support one task. By setting the ‘weight = balanced’ (implemented in Python with sklearn), we tested the performance of three modules on classifying different tasks (one task labeled as ‘1’, and the others labeled as ‘0’).

### Comparison of brain state dynamics between two groups

Independent sample t-tests were applied to study the difference in fractional occupancy and mean lifetime between the stress and control groups for each task and state. FDR correction was used for each task and condition to balance the Type I and Type II errors. We used Network-Based Statistics (NBS, version 1.2) (84) to compare the transition probability in each task. By leveraging the permutation test and family-wise error correction, this method allowed us to test whether some states are more likely to switch to other states in the stress group than the control group and vice versa.

### Permutation testing

To test for differences in correlation effects between stress and control groups, we employed permutation testing. Our approach involved creating 5,000 permutations by shuffling group labels across participants. Subsequently, we performed linear regressions on the two groups, yielding R-values for fractional occupancy and mean lifetime. We then computed the absolute R-value difference between the groups for each permutation, which provided an empirical null distribution of 5,000 differences. To estimate the p-value, we calculated the proportion of null distribution samples that exceeded or matched the between-group R-c in the actual data.

## Supplementary Figures S1-S6

**Figure S1.**
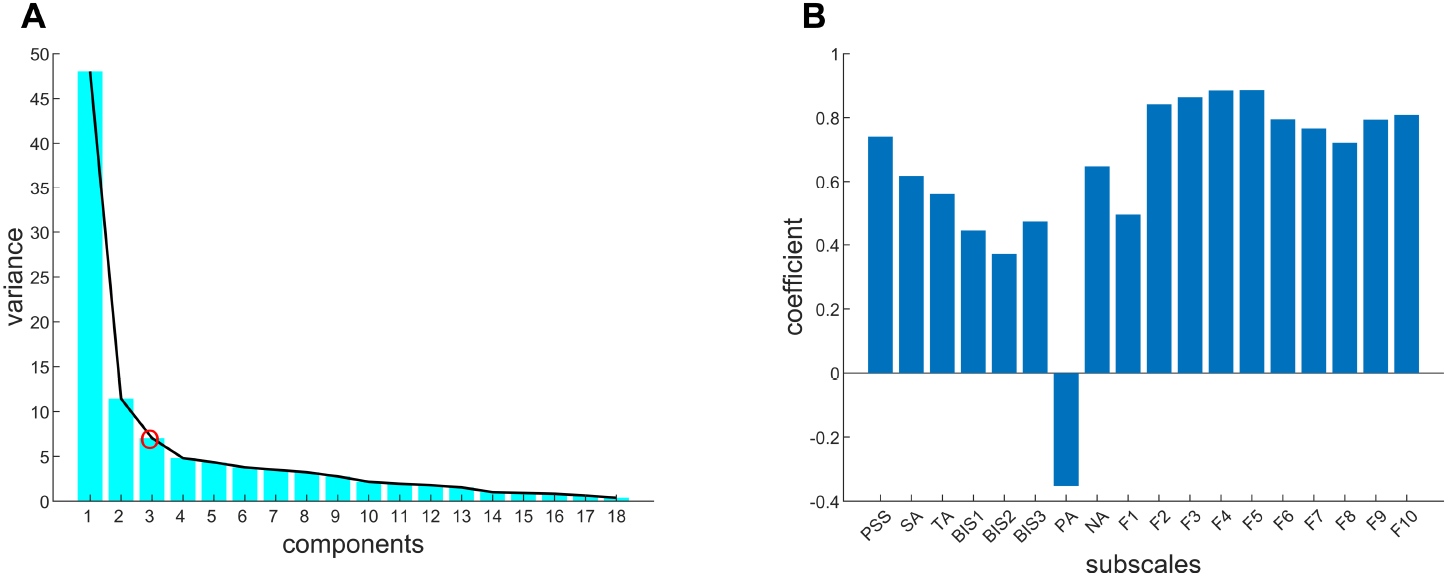
PCA’s results based on psychological assessments. **A**, the bar graph shows the explained variances (Y axis) of each component (X axis), and the red circle represents the last principal component. Therefore, three principal components are extracted from the PCA. **B**, the bar graph shows that the first principal component has positive loadings in all subscales except the positive affect subscale in PANAS (−0.35). Thus, we chose the first principal component as long-term psychological distress. PSS, perceived stress scale. SA, state anxiety. TA, trait anxiety. PA, positive affect. NA, negative affect. BIS, Barratt Impulsiveness Scale. F1-F10, subscales in SCL-90, Symptom Checklist-90.

**Figure S2.**
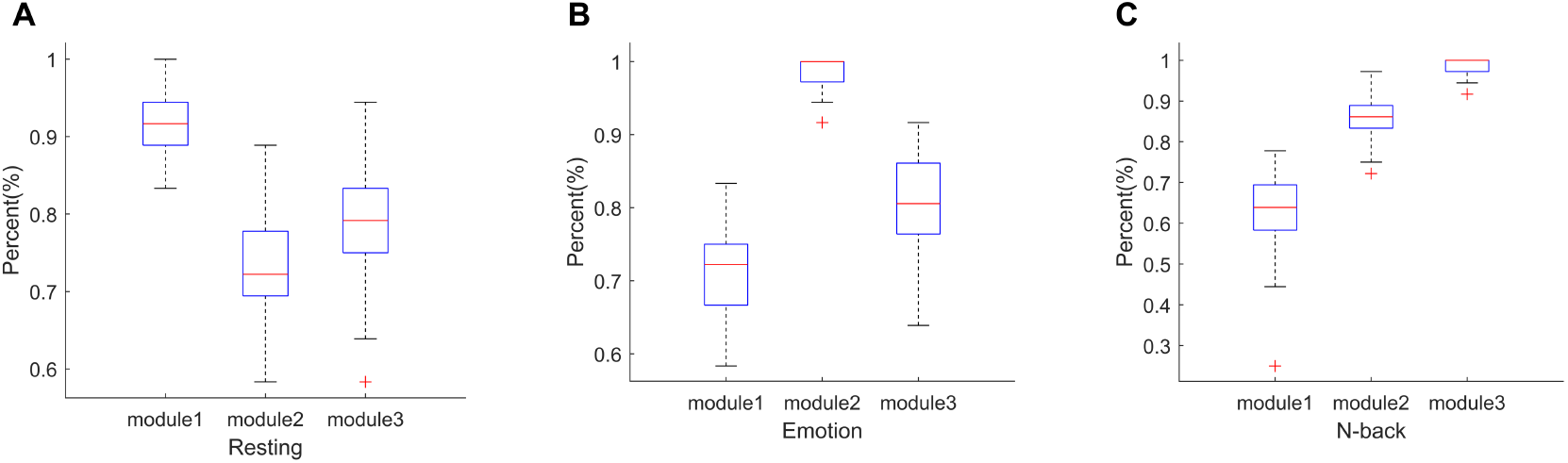
The results of rest, emotional, and working memory processing classification based on transition probability in Module_rest_, Module_emotion_, and Module_WM_. **A-C**, Box plots of accuracy measures from the classification for resting, emotion matching task, and N-back, respectively. During each classification, one was labeled as ‘1’, and the two others were labeled as ‘0’. The transition probability of states in each module was used as task-specific features. Through 100 runs of 5-fold cross-validation on the training data (80%), the accuracy distributions were acquired on the testing data (20%). Notes: Blue box, 25-75 percentile. Red ‘+’, outliers. Module 1, Module_rest_. Module 2, Module_emotion_. Module 3, Module_WM_.

**Figure S3.**
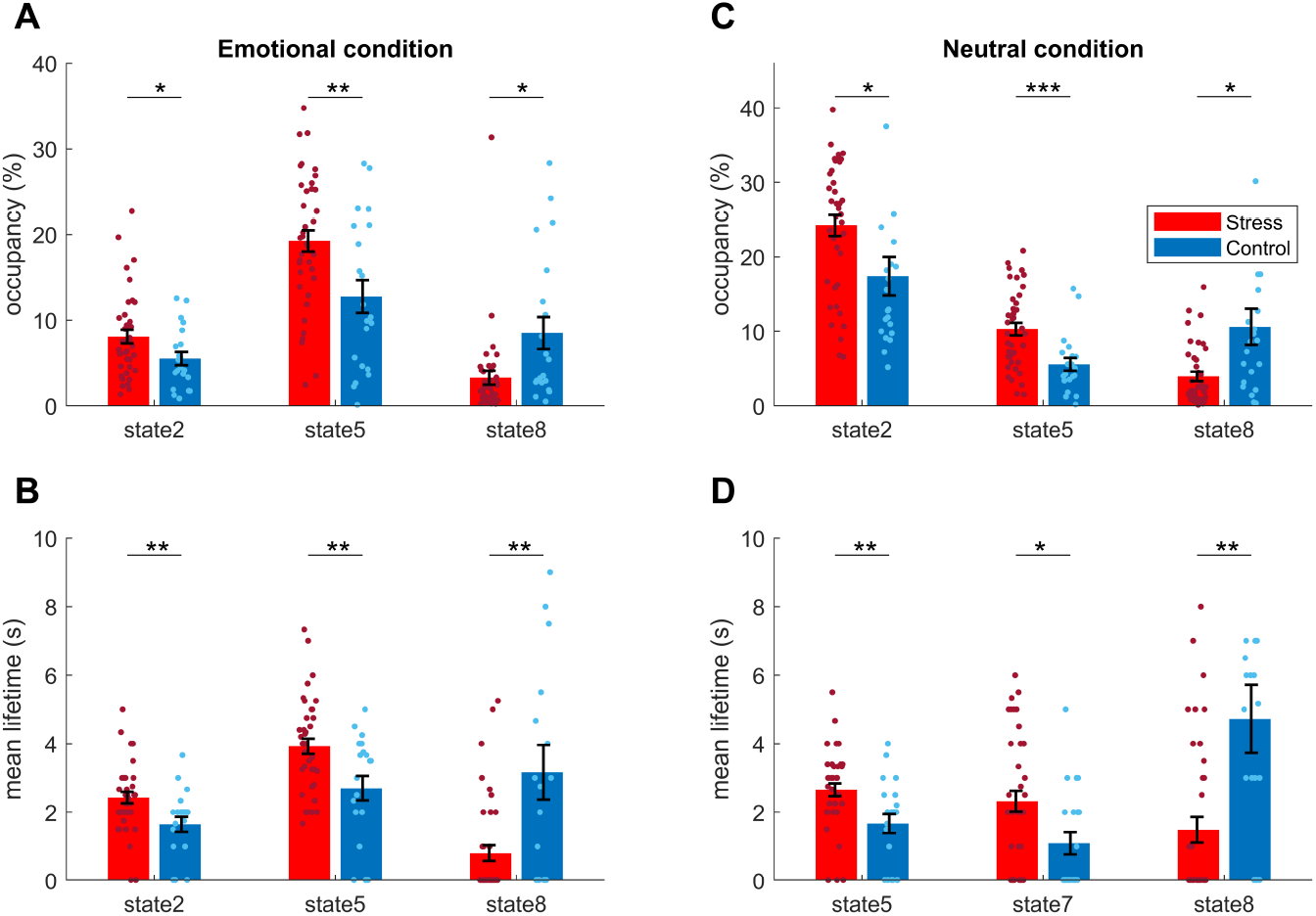
Long-term stress-related changes in brain state dynamics at each condition of emotion matching task. **A-B**, Emotional condition. States 2, 5 (State_emotion_) have higher occupancy in the stress group, whereas state 8 has higher occupancy in the control group. The mean lifetime of states 2 and 5 are higher in the stress group, while the mean lifetime of state 8 is higher in the control group. **C-D**, Neutral condition. States 2, 5 have higher occupancy in the stress group, whereas state 8 has higher occupancy in the control group. The mean lifetime of states 5 and 7 (State_emotion_) is higher in the stress group, while the mean lifetime of state 8 is higher in the control group.

**Figure S4.**
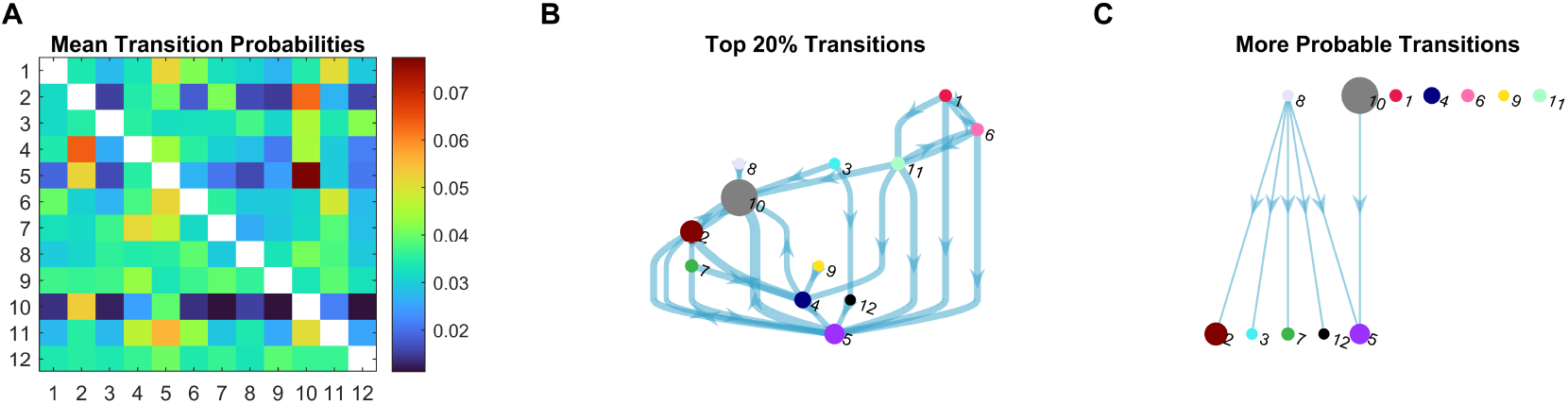
Long-term stress’ effect on the transition probability of emotion matching task. **A**, Group averaged transition probability matrices for the stress group. **B**, The Top 20% transition probability in the stress group. **C**, Transitions are more likely to occur in the stress group than the control group (*P*_FWE_ < 0.05).

**Figure S5.**
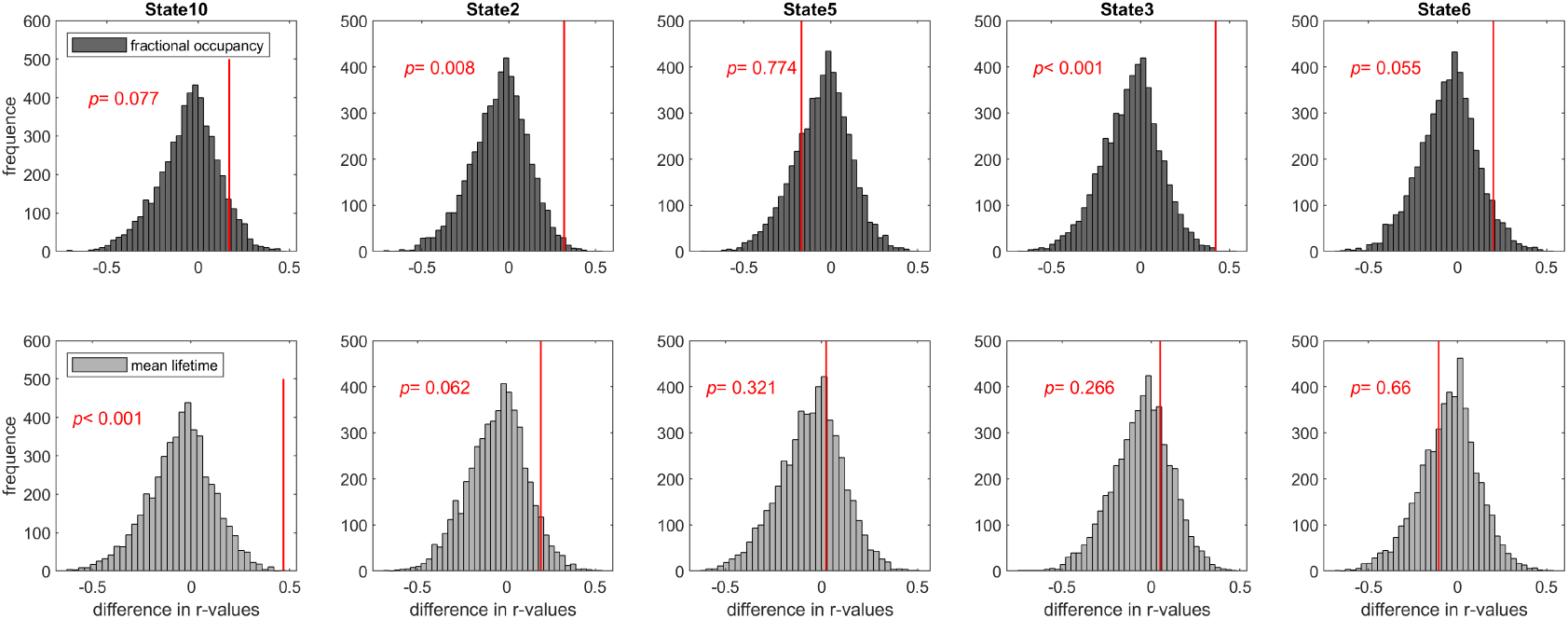
The significant difference in correlation effects between long-term stress and control groups. We created null distribution for each state in fractional occupancy (top panel) and mean lifetime (bottom panel) by generating 5,000 permutations. For each permutation, we used the absolute R-value in the stress group minus the absolute R-value in the control group to calculate the between-group difference. Histograms display the frequency of between-group differences in absolute R-values in permutation data, and red lines represent the real data. P-values were estimated by calculating the proportion of samples in the null distribution that were greater than or equal to the difference of between-group R-values in the real data. Notes: * *P* < 0.05; ** *P* < 0.01; no * 0.05 < *P* < 0.1 (results that fail to pass through the FDR correction were also included in this bar graph).

**Figure S6.**
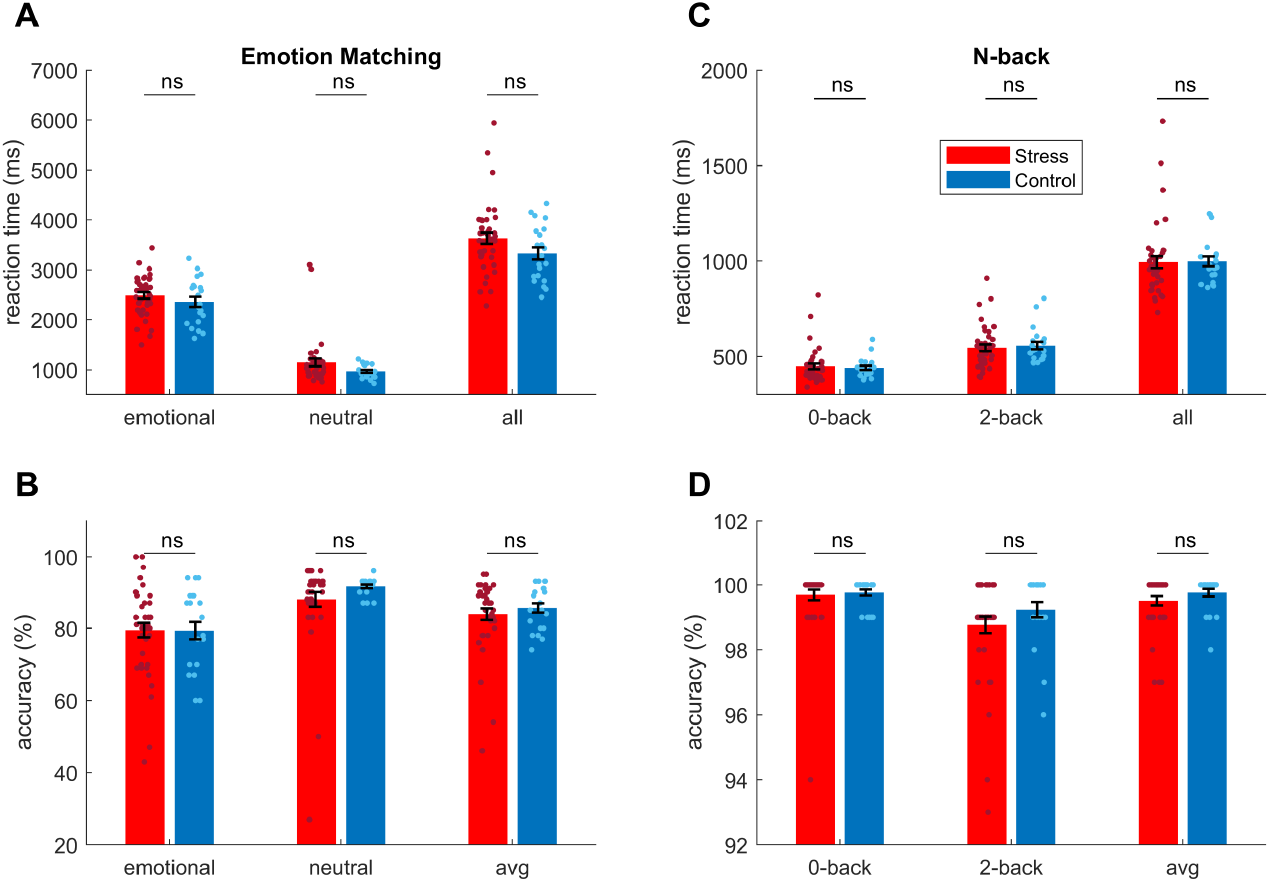
Similar performance in reaction time and accuracy between the long-term stress group and controls. We found that there is no significant difference in reaction time and accuracy of emotion matching task (**A-B**) and N-back **(C-D**) between the stress group and the control group. Emotional, emotional condition in the emotion matching task; Neutral, neutral condition in the emotion matching task. 0-back and 2-back are two conditions in the N-back task. All, the sum of reaction time in two conditions at each task. Avg, the average of accuracy in two conditions at each task. ns, not significant.

**Figure S7.**
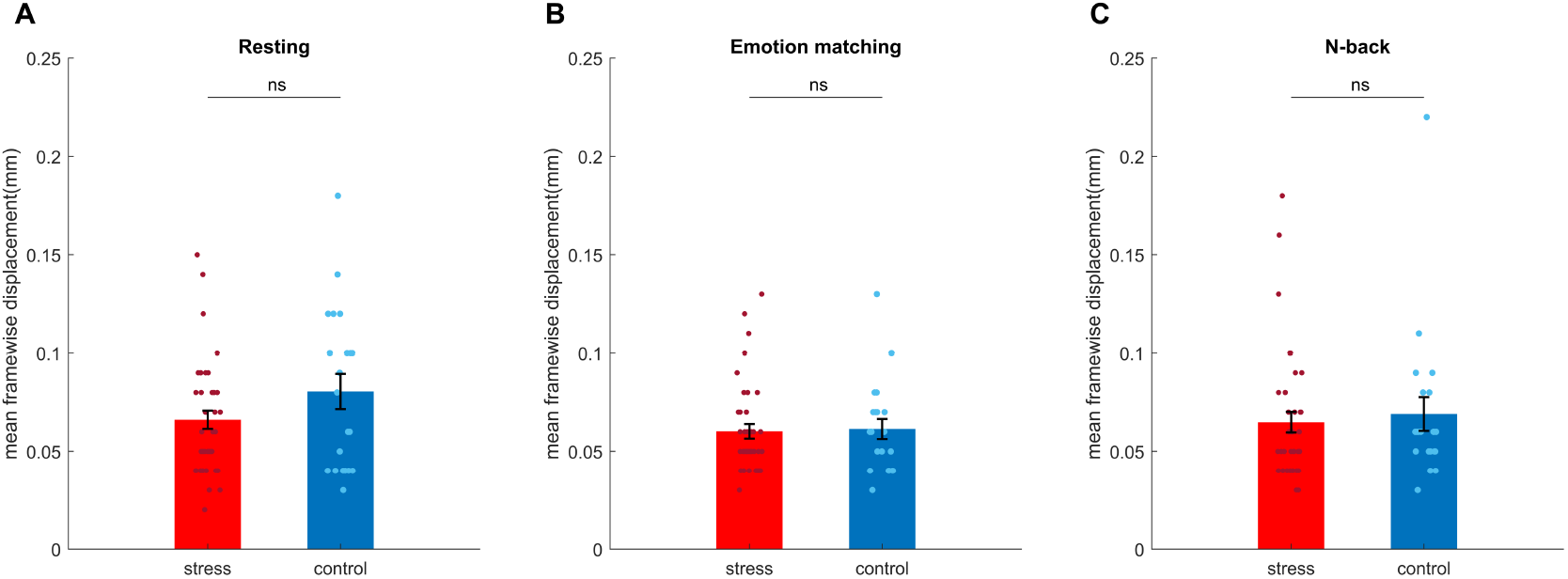
Similar head motion in the scanner between the long-term stress group and controls at each task. There is no significant difference in head motion between the stress and control groups across the resting phase (**A**), emotion matching task (**B**), and N-back (**C**). ns, not significant.

